# Haploinsufficiency of the mouse *Tshz3* gene leads to kidney dysfunction

**DOI:** 10.1101/2021.08.16.456481

**Authors:** Irene Sanchez-Martin, Pedro Magalhães, Ahmed Fatmi, Fabrice Richard, Thien Phong Vu Manh, Andy Saurin, Guylène Feuillet, Colette Denis, Joost P. Schanstra, Petra Zürbig, Xavier Caubit, Laurent Fasano

**Affiliations:** Aix Marseille Univ, CNRS, IBDM, UMR7288, Marseille, France; Mosaiques Diagnostics GmbH, Hannover, Germany; Aix Marseille Univ, CNRS, INSERM, CIML, Centre d’Immunologie de Marseille-Luminy, Marseille, France; Institut National de la Santé et de la Recherche Médicale (INSERM), U1297, Institut of Cardiovascular and Metabolic Disease, Toulouse, France; Université Toulouse III Paul-Sabatier, Toulouse, France

## Abstract

Renal tract defects and autism spectrum disorder (ASD) deficits represent the phenotypic core of the 19q12 deletion syndrome caused by the loss of one copy of the *TSHZ3* gene. While a proportion of *Tshz3* heterozygous (*Tshz3^+/lacZ^*) mice display ureteral defects, no kidney defects have been reported in these mice. The purpose of this study was to characterize the expression of *Tshz3* in adult kidney as well as the renal physiological consequences of embryonic haploinsufficiency of *Tshz3* by analyzing the morphology and function of *Tshz3* heterozygous adult kidney. Here, we described *Tshz3* expression in the smooth muscle and stromal cells lining the renal pelvis, the papilla and glomerular endothelial cells (GEnCs) of the adult kidney. Histological analysis showed that *Tshz3^+/lacZ^* adult kidney had an average of 29% fewer glomeruli than wild type kidney. Transmission electron microscopy (TEM) of *Tshz3^+/lacZ^* glomeruli revealed ultrastructural defects. Compared to wild type, *Tshz3^+/lacZ^* mice showed no difference in their urine parameters but lower blood urea, phosphates, magnesium and potassium at 2 months of age. At the molecular level, transcriptome analysis identified differentially expressed genes related to inflammatory processes in *Tshz3^+/lacZ^* compare to wild type (WT; control) adult kidneys. Lastly, analysis of the urinary peptidome revealed 33 peptides associated with *Tshz3^+/lacZ^* adult mice. These results provide the first evidence that in the mouse *Tshz3* haploinsufficiency leads to cellular, molecular and functional abnormalities in the adult mouse kidney.

## Introduction

Congenital anomalies of the kidneys and urinary tract (CAKUT) are the most common cause of renal failure in children ^1^. Ureteropelvic junction obstruction (UPJO), the most common paediatric renal obstructive disorder, has an incidence of 1 in 1000-1500 live births screened by antenatal ultrasound ^2^. Congenital UPJO is usually caused by the presence of an aperistaltic segment of the ureter, preventing the efficient transport of the urine from the kidney to the bladder. UPJO can result from the decrease in the number of smooth muscle cells, interstitial Cajal-like cells and nerve fibers in the ureteropelvic junction. Therefore, impaired transport of urine can lead to an increase in back-pressure on the kidney, hydronephrosis, and progressive damage to the kidney function ^3^.

The *TSHZ3* gene (Teashirt zinc-finger homeobox family member 3; also known as ZNF537), which encodes a zinc-finger transcription factor, was recently identified as the critical region for a 19q12 deletion syndrome (19q12DS): patients with *TSHZ3* heterozygous deletion show lower (i.e. vesicoureteral reflux grade 2) and upper (i.e. UPJO) urinary tract defects as well as kidney (i.e. hydronephrosis and nephrolithiasis) defects^4^.

*Tshz3* homozygous mutant mice (*Tshz3^lacZ/lacZ^*) have been used to explore the pathogenesis of *Tshz3* in UPJO ^5,6^. Studies have shown that from embryonic day (E) 12.5 onwards TSHZ3 positive cells are detected in the mouse ureter and the kidney ^5^. In the embryo, *Tshz3* plays a key role in smooth muscle differentiation by regulating *Myocardin (myocd)* expression and MYOCD activity ^5,6^. *Tshz3^lacZ/lacZ^* mutant mice die perinatally because of their inability to breathe and display bilateral UPJO and hydronephrosis ^5,7^. In comparison, one-fourth of *Tshz3^+/lacZ^* heterozygous embryos display a unilateral UPJO and hydronephrosis and about 50% die at birth ^5–7^. However, the impact of *Tshz3* haploinsufficiency on the postnatal kidneys has never been investigated so far.

Therefore, the main aim of this study was to determine whether *Tshz3* heterozygote mice exhibit kidney defects in order to gain insights on whether haploinsufficiency may help explaining kidney diseases reported in patients with heterozygous deletion of *TSHZ3*.

## Results

### TSHZ3 expression in adult kidneys

Previous analyses performed during development indicate that the temporal and spatial distribution of β-galactosidase (ß-gal) protein in *Tshz3^+/lacZ^* mice faithfully reproduces the expression of *Tshz3/TSHZ3* ^5,6^. Here we used the same approach to characterize the expression of *Tshz3*/TSHZ3 in sections of *Tshz3^+/lacZ^* adult kidneys. X-Gal staining performed on *Tshz3^+/lacZ^* adult kidney revealed that *Tshz3* is expressed in the pelvic region and ureter, the papilla, perivascular region and the glomeruli (**Fig. 1A-F**). To further characterize the expression of TSHZ3 in the glomeruli, we performed immunostaining, using glomerular cells markers. These analyses showed that β-gal positive cells were endothelial cell (CD31-positive) but not podocytes (Dachshund 1 positive) or mesangial cells (NG2 positive) (**Fig. 1G-I**). Double immunostaining for β-gal and Smooth muscle alpha-actin (SMAA) confirmed the expression of TSHZ3 in smooth muscle cells in the pelvic region and in SMA-negative Dachshund 1-positive stromal cells lining the urothelium (**Fig. 2**).

**Figure 1.**
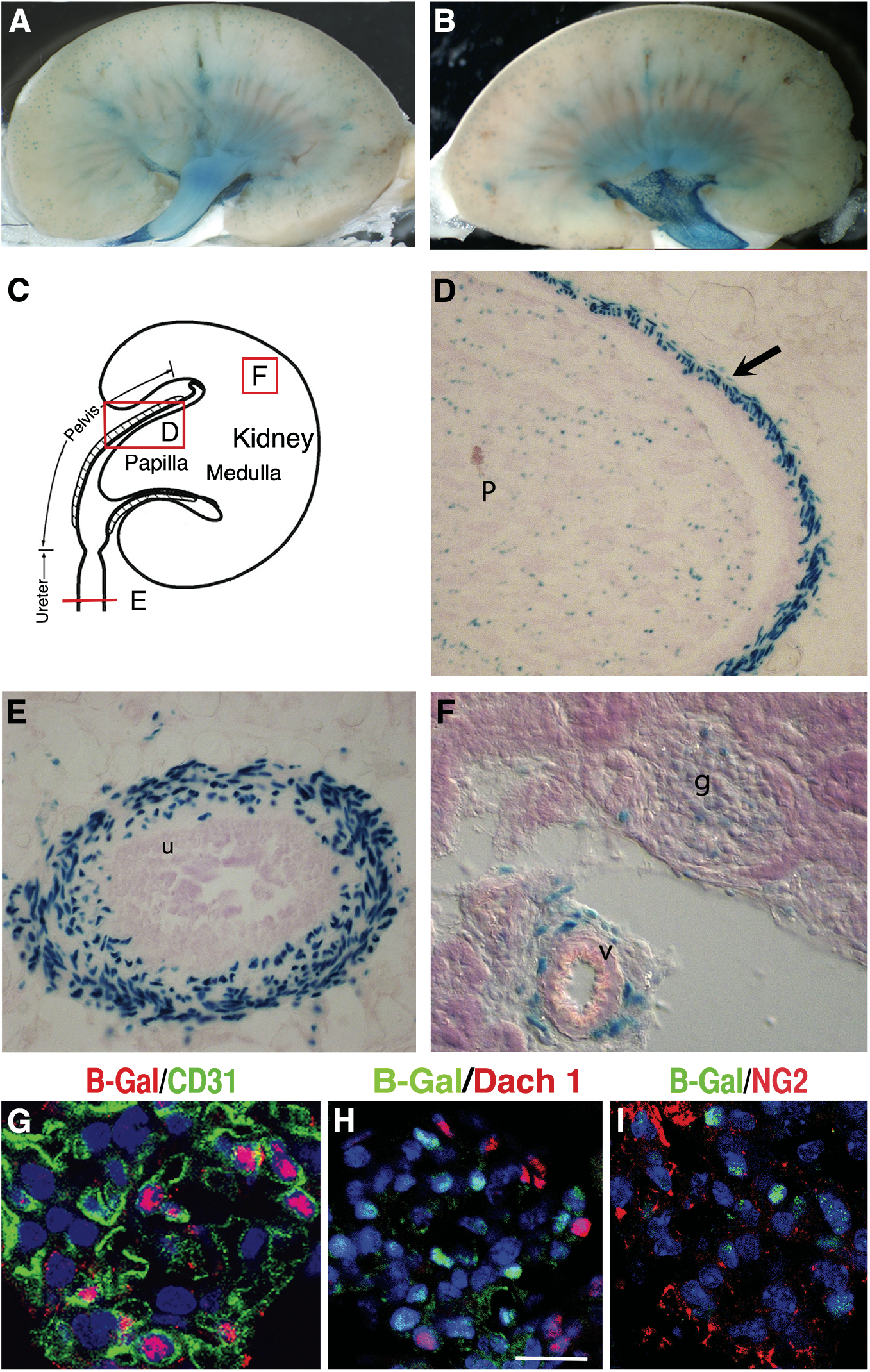
Distribution of Xgal-positive cells in *Tshz3^+/lacZ^* adult renal tract. (A, B) The two halves of the same adult kidney stained with X-Gal. (C) Cartoon showing the section sites and planes for D, E and F. (D-F) Xgal-Positive cells were found in papilla (p) and smooth muscle present in the pelvic region (arrow) (D), in the mesenchymal part of the ureter (E), in the glomeruli (g) and in close proximity to blood vessel (v) (F). (G-I) In glomeruli, TSHZ3 (ß-Gal) is detected in CD31+ glomerular endothelial cells (G) but not in DACH1+ podocytes (H) or in NG2+ mesangial cells (I); scale bar, in G-I, 20 μm. B-Gal, Beta-Galactosidase; u: urothelium.

**Figure 2.**
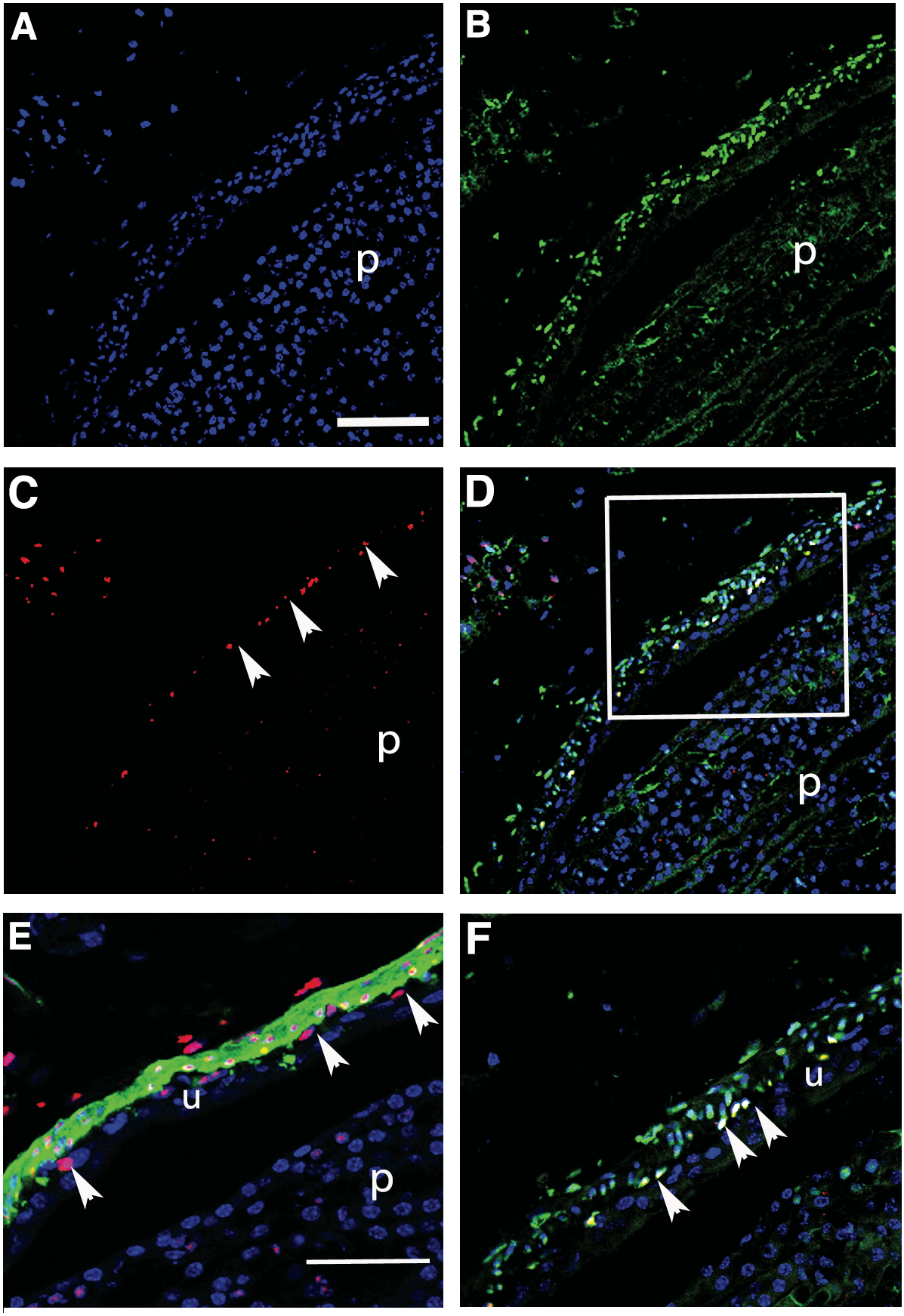
Characterization of β-galactosidase-positive cells in the pelvic region of *Tshz3^+/lacZ^* adult kidney. (A-D) Comparative expression of ß-galactosidase (green) and Dachshund 1 (red) in the pelvic region. (F) Close up of the region boxed in D. Merge image (D, F) shows that a subset of βgal+ cells expresses DACH1 (arrowheads in C and F). These cells represent a cell layer in a sub urothelial position. (E) Double immunostaining for βgalactosidase (red) and smooth muscle actin (green) indicates that βgal expression is found in SMA expressing cells. Some βgal+ cells adjacent to the urothelium do not express SMA (arrowheads in E). P: papilla, u: urothelium. Scale bars, in A, 100μm; in E, 50μm.

### *Tshz3^+/lacZ^* mice showed glomerular defects

Comparison of 8-week-old adult kidney histology revealed a decrease of 28.8% in glomerular density in *Tshz3^+/lacZ^* compared to WT (**Fig. 3A, B and supplementary Fig. 4**). The same analysis performed with 40-week-old adult kidneys revealed a 44.2% reduction of glomerular density in *Tshz3^+/lacZ^ vs*. WT. In order to identify glomerular morphological changes in *Tshz3^+/lacZ^* mice, we conducted an ultrastructural analysis using transmission electron microscopy (TEM). While this analysis did not identify a significant variation of the proportion of the size of the fenestration and endothelial layer (**Fig. 3C, D**), it revealed a significant reduction of the thickness of the glomerular basement membrane (GBM) in glomeruli of *Tshz3^+/lacZ^* mutants (144.2 ± 4.2 nm) compared to WT (155.1 ± 3.32 nm) (**Fig. 3E, F**). TEM analysis also showed a significantly increased foot process width (351.7 ± 5.03 nm) in *Tshz3^+/lacZ^* compared to WT (303.4 ± 6.57 nm) podocytes, suggesting foot process effacement (FPE) (**Fig. 3G, H**).

**Figure 3:**
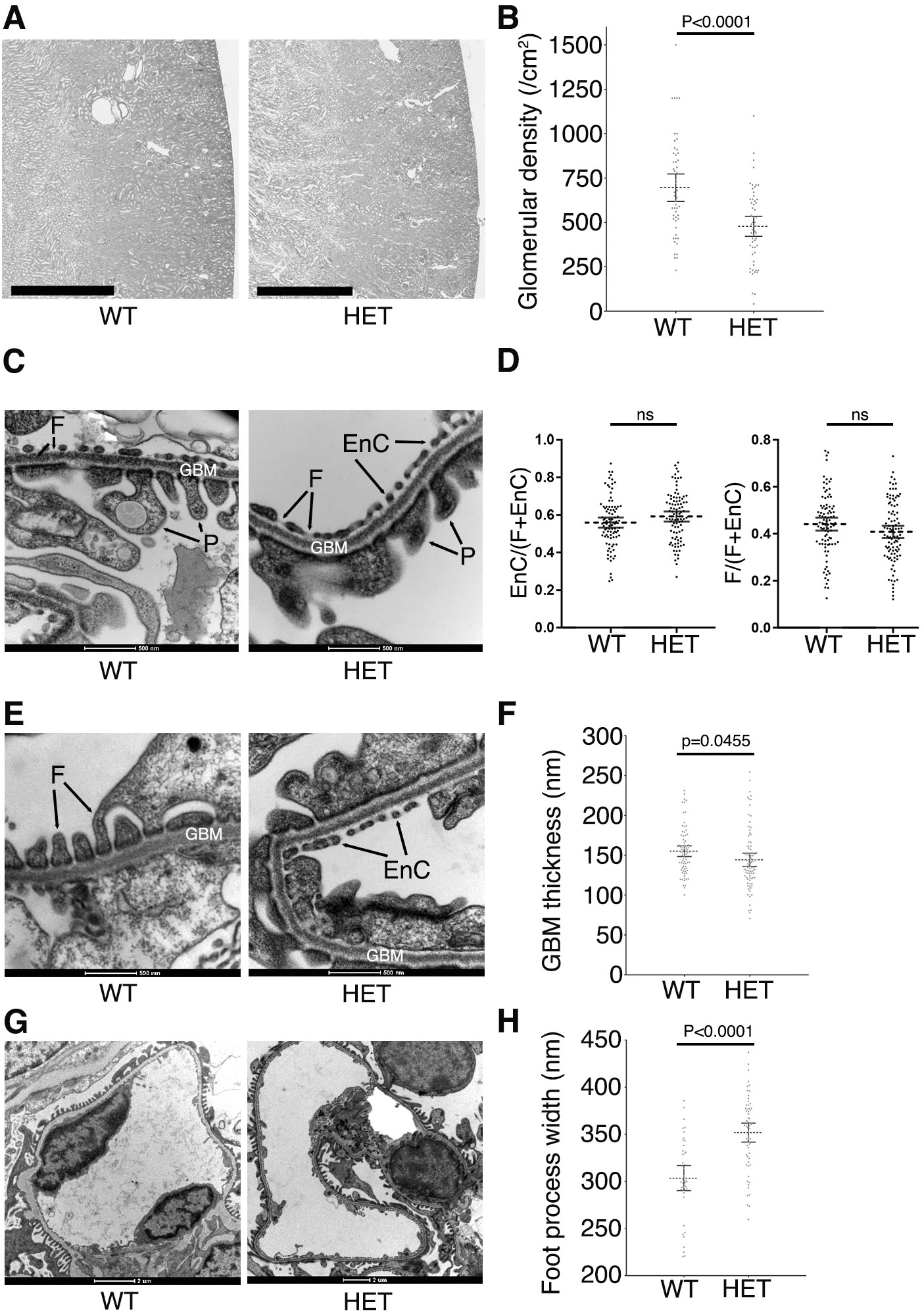
Reduced glomerular density, glomerular basement membrane thinning and foot process effacement in heterozygous *Tshz3^lacZ/+^* mice. (A) Representative images of haematoxylin and eosin-stained sections of WT and *Tshz3^+/lacZ^* adult kidneys at post-natal day 60. Scale bar 1mm. (B) The graph shows the significant (P<0.0001) reduction of the glomerular density in heterozygous *Tshz3^lacZ/+^* (471 ± 0.29 /cm^2^, 48 sections from 3 kidneys) compare to wild-type kidney (695 ± 0.38 /cm^2^, 41 sections from 3 kidneys). (C, E, G) Representative TEM images of wild type and *Tshz3^+/lacZ^* adult kidneys. (D, F, H) TEM morphometric analysis. (C) TEM: No abnormalities in endothelial cell fenestration are observe in *Tshz3^+/lacZ^* mice. Scale bar, 500nm. (D) Morphometric analysis of the fenestration reveals no significant difference between WT and *Tshz3^+/lacZ^* mice. (E) TEM: *Tshz3^+/lacZ^* kidney shows a reduction of the thickness of the GBM in *Tshz3^lacZ/+^* compared to WT. Scale bar, 500nm. (F) Morphometric analysis reveals a significant reduction (P<0.0455) of the thickness of the GBM in *Tshz3^lacZ/+^* (144.2 ± 4.2 nm, 84 sections from 3 kidneys) compare to wild-type (155.1 ± 3.32 nm, 78 sections from 3 kidneys). (G) TEM: illustrating increased foot process effacement in *Tshz3^lacZ/+^*compared to WT mice. Scale bar, 2μm. (H) Foot process width in WT (303.4 ± 6.57 nm, 42 sections from 7 kidneys) is significantly lower (P<0.0001) compared to *Tshz3^lacZ/+^* (351.7 ± 5.03 nm, 61 sections from 9 kidneys) mice. Data are shown as mean and its 95% CI. EnC, endothelial cell cytoplasm; F, endothelial fenestration; GBM, glomerular basement membrane; HET, hererozygous; P, podocyte foot process; SD, standard deviation; WT, wild type.

### Blood electrolytes are modified in *Tshz3^+/lacZ^* mice

To characterize the effects of *Tshz3* haploinsufficiency on kidney filtration, we performed biochemical measurements on blood samples and generated biochemical profiles for *Tshz3^+/lacZ^* and WT adult mice tested at 58-64 days-of-age. While this analysis demonstrated that *Tshz3^+/lacZ^* mice had no proteinuria compared to WT, it showed a significant reduction of the urea (P<0.013), phosphates (P<0.011), magnesium (P<0.014), potassium (P<0.0034) as well as an increased concentration of sodium (*P* <0.012) (**Fig. 4; Table 1a**). This analysis also suggested a trend for an increased concentration of creatinine in *Tshz3^+/lacZ^* compared to WT (15.91 ± 1.51 vs. 12.36 ± 2.07 μm/l), but no statistical significance was observed.

**Figure 4.**
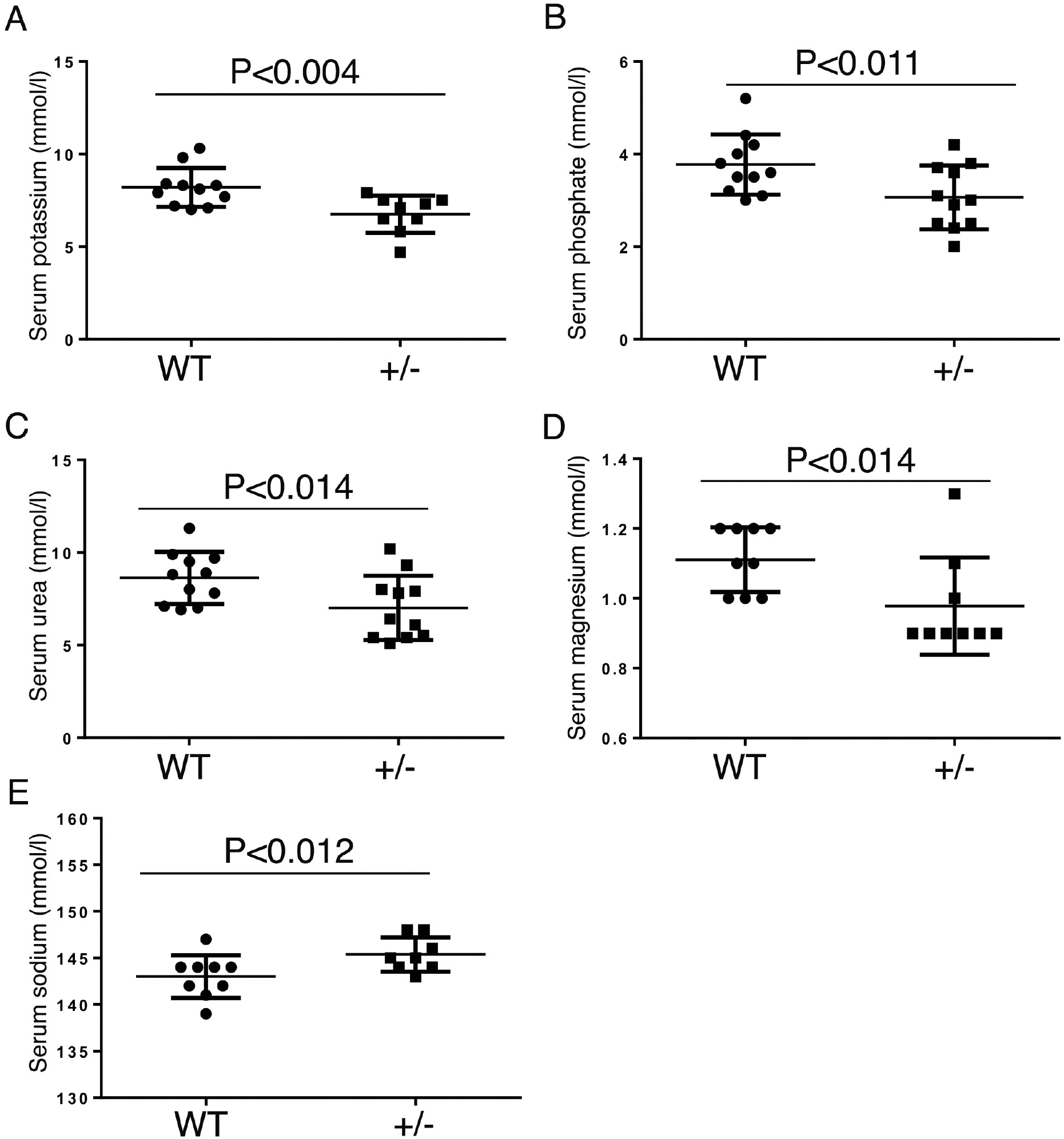
Differences in serum biochemical parameters between control and *Tshz3^lacZ/+^* mice. Plasma concentration of potassium (A), phosphates (B), urea (C), magnesium (D) and sodium (E). Data are represented as means ± SEM. Statistically significant difference from control at *p<0.02; **p<0.004. WT, wild type; +/-, *Tshz3^+/lacZ^*

### The *Tshz3^+/lacZ^* adult kidney showed differential expression of genes involved in inflammatory processes

To identify *Tshz3*-regulated genes in the adult kidney, RNA sequencing analysis was performed with samples extracted from *Tshz3^+/lacZ^* (n=6) and wild type (WT; controls, n=6) adult mouse kidneys. 97 differentially expressed genes (DEGs) were identified, among which 55 were up-regulated and 42 were down-regulated (adjusted p-value < 0.01) (**Table 1b**). Thereafter, we sought to take advantage of the single-cell transcriptome of mouse glomeruli ^8^ and adult kidney^9^ to identify the cell types in wild type kidneys that expressed the genes differentially expressed in *Tshz3^+/lacZ^* kidney. This analysis identified 58 genes expressed in podocytes, 31 in mesangial cells, 27 in immune cells, 40 in tubules and 43 in GECs, which might be direct targets of TSHZ3 (**Table 1c**). We also found that some DEGs were shown to be markers of proximal tubules (*Aspdh*, *Snhg11* and *D630029K05Rik*, a long non-coding RNA) or myeloid lineage (*Cd74*, *Gusb*, *H2-Aa*, *H2-Ab1*, *Lyz2*, *Mpeg1* and *Vcam1*), including macrophages and dendritic cells ^10^.

To characterize the differential expression data and to identify biological changes that occur in *Tshz3^+/lacZ^* kidneys, we performed different enrichment analyses. As only a few genes showed very strong differential expression, we applied gene set enrichment analysis (GSEA ^11,12^) using a ranked gene list. We chose the updated public genesets available on MSigDB ^12^ and identified positive enrichment in the kidneys of *Tshz3^+/lacZ^*adult mice for gene sets related to “interferon-gamma response”, “epithelial to mesenchymal transition”, “inflammatory response” and negative enrichment for genes related to “Oxidative Phosphorylation” and “Xenobiotic Metabolism” (**Table 1d; Supplementary Fig. 1**). Furthermore, ingenuity pathway analysis (IPA) of the DEGs revealed enrichment of cellular processes centered on inflammatory and kidney diseases (**Supplementary Fig. 2A, B**). This analysis showed that the relevant toxicity phenotypes and pathology endpoints associated with the DEGs in *Tshz3^+/lacZ^* mice were centered on the kidney and identified interferon-gamma (IFNG) as an upstream regulator (**Supplementary Fig. 2B, C**). Moreover, enrichment analysis of pathways and transcription factor using enrichR ^13^ also identified the *“interferon gamma”* pathway but also *“collagen formation”* and *“extracellular matrix organization”* as significantly enriched in *Tshz3^+/lacZ^* adult kidneys (**Table 1e**). Interestingly, the ChIP enrichment analysis (ChEA) and a database search revealed that 22 DEGs (21.68%) are direct targets of the transcription factor interferon regulatory factor 8 (IRF8) (**Table 1f**).

Using RT-qPCR we confirmed the DEG status observed by RNA-seq analysis for *Ciita*, *Pld4* (two targets of IRF8), *Tlr7* (IRF8 target that promotes IFNG production ^14^) and the proinflammatory *Npy* (**Supplementary Fig. 3**).

### Genes differentially expressed in *Tshz3^+/lacZ^* adult kidney are associated with ASD

To gain insights into *TSHZ3* function, we studied the disease association of the 80 non-ambiguous human orthologs of the mouse DEGs (**Table 1g**). This analysis identified 35/80 genes (43.75%), which were established or putative causes for kidney disorders. Interestingly, 8 of these genes have been also associated with ASD (**Table 1h**).

### Identification of *Tshz3^+/LacZ^*-related urinary peptides

Since *Tshz3^+/lacZ^* mice exhibit dysregulated kidney expression of several genes and altered hematological parameters, we sought to analyze their urinary peptidome. Urine samples derived from *Tshz3^+/LacZ^* (n=15) and WT (n=12) were analyzed by capillary electrophoresis coupled to mass spectrometry (CE-MS). Comparison of urinary peptidome profiles led to the identification of 33 peptides that were significantly (adjusted p-value < 0.05) associated with *Tshz3^+/lacZ^* (**Fig. 5**). Protein fragments from clusterin (CLUS), complement factor D (CFAD), histone H2B type 1-F/J/L (H2B1F), major urinary protein 17 (MUP17), tripeptidyl-peptidase 1 (TPP1) and uromodulin (UMOD), as well as a large number (27) of collagen fragments, were identified (**Table 1i)**. *Tshz3^+/lacZ^* mice showed a decreased concentration of uromodulin and an increased concentration of peptides from clusterin, complement factor D, histone H2B type 1-F/J/L, major urinary protein 17 as well as tripeptidyl-peptidase 1 fragments. Twelve collagen peptides were in higher abundance and fifteen in lower abundance. As UPJO can lead to significant kidney damage, we compared the identified 33 urinary peptides from mice with those of the CKD273 classifier, a predictive marker of chronic kidney disease (CKD) progression in humans^15^. This analysis identified 14 similar (orthologs) human peptides in the CKD273 classifier, most of which were collagen fragments (10 from collagen type I alpha-1 chain and 3 from type III alpha-1 chain). One peptide fragment was from uromodulin.

**Figure 5.**
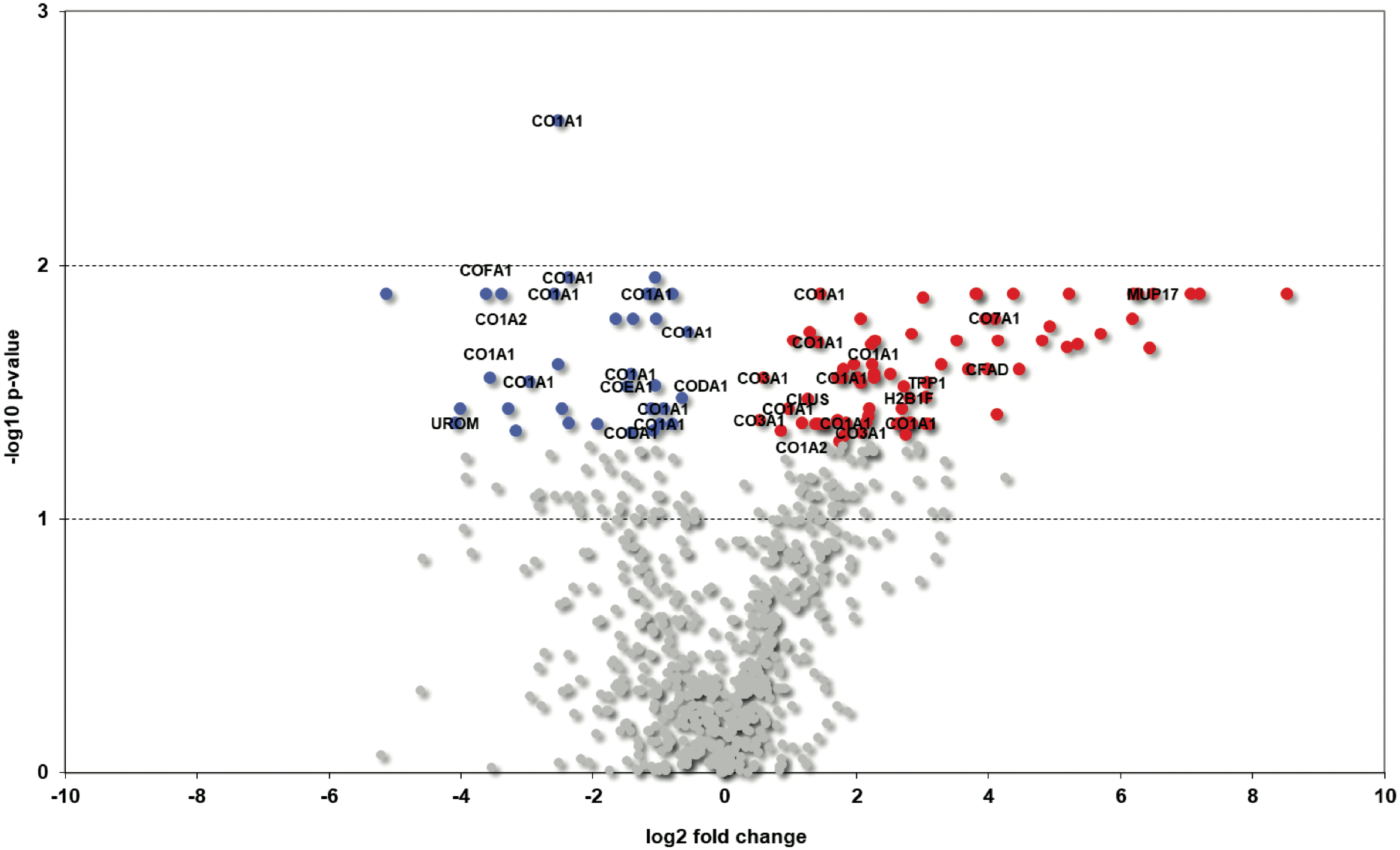
Urinary peptides displaying differential abundance in *Tshz3^+/LacZ^* mice. The volcano plot of quantified proteins revealed that 115 peptides (red and blue dots) show a significant change (p-value <0.05) in *Tshz3^+/lacZ^* compared to control mice. However, for 33 of these peptides we were able to find a sequence (labeled ones; see **Table 1i**). The grey dots are peptides that have not a significant change (p-value >0.05).

## Discussion

Previously, we reported that in mice and humans, haploinsufficiency of *Tshz3/TSHZ3* results in hydroureter ^5,16^ and that individuals with *TSHZ3* heterozygous deletion are at risk to develop kidney diseases ^16^. However, the expression of *Tshz3* in adult kidney and the morphology of *Tshz3* heterozygous adult kidney have not been investigated before. Herein, we characterized TSHZ3 expression in the adult mouse kidney, in particular in glomeruli where TSHZ3 is specifically expressed in endothelial cells (GEnCs), which is supported by single-cell RNA sequencing analysis ^8^. Using *Tshz3^+/lacZ^* heterozygous mice, we have also uncovered a key role for the TSHZ3 transcription factor in controlling the glomerular density and morphology. Notably, we showed that *Tshz3^+/lacZ^* mice have abnormal blood electrolytes and identified *Tshz3^+/LacZ^*-related urinary peptides. In addition, by coupling transcriptomics with *in silico* analysis, we showed that in kidneys from *Tshz3^+/LacZ^* mice, the expression of genes associated with inflammatory processes and of renal- and ASD-associated was different from that in WT mice. Combined with our previous reports, our data suggest that the *Tshz3* heterozygous mice constitute a model that replicates many of the corresponding human disease phenotypes.

The GEnCs, together with the glomerular basement membrane (GBM) and the podocytes, constitute the glomerular filtration barrier (GFB), which selectively filtrates the plasma. The GBM derives from the fusion of the basement membranes of both GEnCs and podocytes, and in adult kidney, the GEnCs may contribute to the renewed biosynthesis of the GBM. Because abnormalities in each of the three constituents of the GFB can lead to proteinuria and kidney disease, TEM was used to assess fine structures of the GFB. While TEM analysis revealed no obvious structural phenotype in TSHZ3-expressing GEnCs, it provided evidence that *Tshz3* haploinsufficiency leads to GBM thinning and change in podocyte morphology evidenced by foot process effacement (FPE). Interestingly, individuals with a thin GBM have hematuria but minimal proteinuria ^17^ and, accordingly, we found that the thin GBM as well as the defects in the podocytes did not result in the development of proteinuria. In future research it would be interesting to investigate whether these defects are associated with hematuria. At present, the primary cause of the structural defects observed in the GBM and the podocytes remains unknown. However, since communication between intraglomerular cells is required for proper development and maintenance of the GFB ^18^, GBM thinning and foot process effacement might be the direct or indirect consequences of an abnormal cross-talk between the TSHZ3-positive GEnCs and other glomerular cells. Furthermore, *Tshz3* may be transiently expressed in glomerular cell lineages. Nevertheless, the comparison of the urinary peptidome of *Tshz3^+/lacZ^* and WT adult led to the identification of 27 collagen and 6 non-collagen peptides associated with *Tshz3^+/lacZ^* mice. Among the collagen fragments displaying a deregulation (e.g. 12 increased or 15 decreased), we identified thirteen collagen fragments that overlap the CKD273 classifier that might be indicative of the development of fibrosis ^19,20^. As previously suggested ^21^, the fragments overlapping the human CKD273 classifier could be used as biomarkers to assess renal function in patients with *TSHZ3* heterozygous deletion. Interestingly, our analysis of urinary peptides also revealed an under-representation of a MUP17 peptide in *Tshz3^+/lacZ^* mice. The MUP17 protein is predominantly expressed in males and dominant males significantly increased the secretion of MUP17 in social conditions ^22,23^. In the future, it would be interesting to study the relationship between variations in the level of MUP17 and the severity of the social behavior deficit observed in *Tshz3^+/lacZ^* mice. To further evaluate renal function, we performed biochemical measurements on blood samples and generated biochemistry profiles. This analysis identified a significantly reduced concentration for urea, phosphates, magnesium and potassium in *Tshz3^+/lacZ^* mice. Reduced serum urea is less frequent and usually of less clinical significance than increased serum urea ^24,25^. Nevertheless, overhydration is one of the rare causes of decreased serum urea and we routinely observed that *Tshz3^+/lacZ^* mice drink more and urinate more than WT mice (not shown). The lower plasma magnesium (hypomagnesaemia), phosphate (hypophosphatemia) and potassium (hypokalemia) serum concentration are associated with defective reabsorption/excretion process in the distal nephron ^26–28^. Interestingly, single-cell transcriptional profiling of the healthy mouse kidney showed expression of *Tshz3* in the proximal tubule, distal convoluted tubule as well as in collecting duct principal and intercalated cells ^9^, suggesting that *Tshz3* haploinsufficiency might impact different components of the nephron. In the future, these transcriptomic data should be complemented by a comparative analysis of the expression of TSHZ3 and segment-specific tubular markers in WT and *Tshz3^+/lacZ^* adult kidneys. To identify pathways that might be altered, we performed RNA-seq analysis and detected 48 statistically significant changes in gene expression in adult kidneys of *Tshz3^+/lacZ^* as compared to WT. Based on the results of enrichment analyses, we detected the involvement of inflammation-related pathways such as interferon-gamma. Of note, seven DEGs (*Crym*, *Ctss*, *Fasn*, *Fcgrt*, *Gabrb3*, *Lyz2*, and *Vcam1*) were found to be associated with kidney diseases, including renal inflammation, crescentic glomerulonephritis or end-stage renal failure. Interestingly, population-based studies support that autism spectrum disorder (ASD) and kidney disease coexist in several genetic disorders (for review see Table 2 in ^29^), suggesting that the same genetic modification can affect neurodevelopment and nephrogenesis. However, because the genetic alterations (deletion or duplication) associated with these disorders often encompass several genes, it is still unclear how these genes contribute to the underlying molecular mechanisms. In this context, the *TSHZ3* gene which associates ASD with a congenital kidney condition is quite unique^4^ and our findings that heterozygous deletion of *Tshz3* alters some of the key functions performed by the kidneys might be relevant for patients with *TSHZ3* heterozygous deletion. Indeed, we previously reported heterozygous deletion of the *TSHZ3* gene in six patients with renal tract defects, including one with nephrolithiasis and a second one with postnatal echogenic kidney and low glomerular filtration rate (Table1 in ^4^) who required renal transplant (Table 2 in ^30^). These results might also provide to be of interest for cognitive defects link to TSHZ3 haploinsufficiency. For now, the role of low plasma concentration of magnesium in ASD is still a matter of debate. While two studies did not find a statistically significant difference in levels of magnesium in children diagnosed ASD ^31,32^, two other studies ^33,34^ demonstrated lower levels of magnesium in children diagnosed with ASD. So far, *Tshz3* mouse models have been shown to be quite analogous to the clinical problems (i.e. ASD-associated deficits, hydroureter and hydronephrosis) reported in patients with *TSHZ3* heterozygous conditions ^4,5^. Our results generate new hypotheses that might lead to further understanding of the clinical problems and to a better diagnosis management of *TSHZ3* patients.

## Materials and methods

### Mouse strain and genotype

The *Tshz3^+/LacZ^* mouse line has been described previously ^5^.

### Samples collection and RNAseq

For RNA sequencing, *Tshz3^+/LacZ^* and wild-type mice kidneys were collected at 60 days-of-age. The mean values for body weight of the *Tshz3^+/LacZ^* and wild-type were 34.7± 3.9 g (n=6) and 34.6 ± 6.5 g (n=6), respectively. Dissected kidneys were stored in RNAlater solution (Qiagen) and kept frozen at -80 °C until RNA extraction.

Total individual kidney RNA was extracted using anRNeasy Maxi kit75162 Lot.260018727/ Lot.160012031 from Quiagen according to manufacturer’s instructions. The integrity of RNA was assessed using a chip-based capillary electrophoresis machine and RNA concentration determined using a full spectrum (220-750nm) spectrophotometer. The quality control of the RNA was additionally checked with RNA 6000 Pico de Agilent Technologies, according to the manufacturer’s instructions. To obtain two independent total RNA preparations from the two different conditions (wild type: WT1 & WT2; *Tshz3^+/LacZ^*: HET1 & HET2) we pooled RNA from 3 kidneys per group in the same proportion. The starting material (1μg Total mRNA, dissolved in RNase-, DNase- and protease-free molecular grade water) was sent to GATC (Eurofins) for sequencing (Genome Sequencer Illumina HiSeq).

Sequences (fastq format) were mapped to the mm10 version of the mouse genome to generate Sequence Alignment/Map (SAM/BAM) format. After normalization, analysis of differentially expressed genes (DEGs) was performed using both the Bioconductor (http://www.bioconductor.org) package DESeq/DESeq2 and the package edgeR.

This analysis generated differential expression lists with False Discovery Rates (FDRs), which are derived from p-values corrected for multiple testing using the Benjamini-Hochberg method. 6 files in total were generated: FDR 1%, 10% for both UCSC (transcripts) and ENSEMBL (genes).

EnrichR tool ^13^ was used to performed enrichment analysis with “pathways”, gene ontology biological processes (GOBP) and transcription factor (ChIP Enrichment Analysis, ChEA). Gene set enrichment analysis (GSEA) was performed using the software provided by the Broad Institute ^35,36^ with default parameters and a pre-ranked gene list calculated based on the10 negative log10 of the *P*-value from DESeq2 analysis multiplied by the sign of differential expression.

### mRNA extraction, cDNA synthesis and quantitative real-time PCR (RT-qPCR)

Total RNA from adult kidneys of WT and *Tshz3^+/lacZ^* mice was prepared using a RNeasy Maxi Kit (ref 75162 Qiagen^™^), and first-strand cDNA was synthesized using a Maxima First Strand cDNA Synthesis Kit with dsDNase (ThermoFisher Scientific^™^ ref K1671 50). All samples from each experiment were reverse-transcribed at the same time, and real-time PCR was performed on a StepOne+ qPCR detection system (Applied Biosystems^™^) using Luminaris Color HiGreen High ROX qPCR Master Mix (Thermo Fisher Scientific^™^ ref K0362). RT-qPCR conditions were as follows: 40 cycles of 95 °C for 15 s and 60 °C for 60 s. Reactions were run in triplicate in three independent experiments. The geometric mean of the housekeeping gene GAPDH was used as an internal control to normalize variability in expression levels, and samples were also normalized to their respective control group. Specificity of reactions was verified by melt curve analysis. Primer sequences used for Sybr qPCR are as follows:

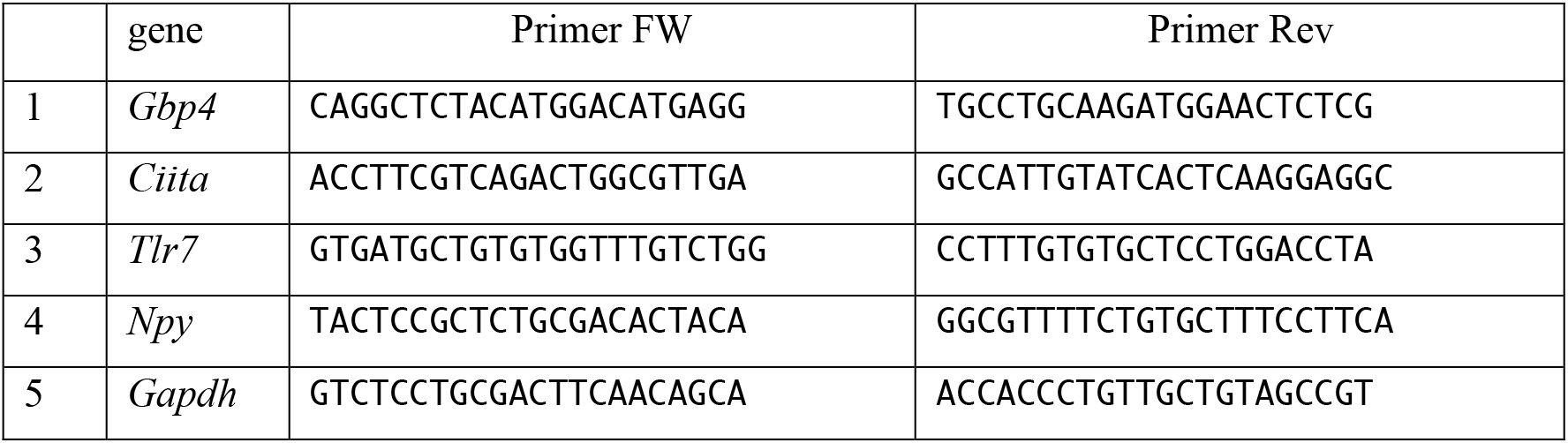

Expression data were normalized to controls and the variability in expression levels was analyzed using the 2^-ΔΔCT^ method described Livak and Schmittgen ^37^.

Variables which showed a p-value less than 0.05 in the resulting model were considered to have a significant effect. These statistical analyses were performed by unpaired t-tests with the qbase+ software version 2 (Biogazelle).

### Immunological and in situ hybridization analysis

Adult mice were transcardially perfused with phosphate-buffered saline (PBS, 10 mM, pH 7.4), followed by 4% paraformaldehyde (PFA) in PBS under ketamine (150 mg/kg) and xylazine (20 mg/kg) anesthesia. Kidneys were post-fixed in 4% paraformaldehyde (PFA; EMS Lot.160401) for 2 h at 4 °C, cryoprotected with 30% sucrose solution in PBS and frozen (Iso-Pentane RPE524391 Carlo Erba at dry ice temperature). Immunostaining was performed on 12-μm cryosections from tissues embedded in OCT compound. Cryosections were washed with 0.2% Tween/PBS for 15 min and then blocked in 10% goat serum 0.1%/0.1% Tween/PBS for 1 h. Sections were incubated with primary antibodies overnight at 4 °C. Secondary antibodies were incubated 2 h at room temperature, and after several washes, sections were counterstained with DAPI for 10 minutes.

The following primary antibodies were used: anti-β-galactosidase (chicken, 1/1000, ab9361, Abcam; rabbit, 1/1000, Cappel), mouse anti-NG2 (1/100, MAB5384 Millipore), rabbit anti-CD31/Pecam (1/100 MEC13.3, BD Pharmingen) and rabbit anti-DACH11 (1/100, Proteintech 10914-1-AP). Secondary antibodies: Alexa-Fluor-546 goat anti-mouse; Alexa-Fluor-546 donkey anti-rabbit, Alexa-Fluor 488 donkey anti-mouse, Alexa-Fluor-488 donkey anti-chicken. Slides were mounted with Fluoromount-G, Southern Biotech Lot: B0216-N156. Images were acquired using a laser scanning confocal microscope (LSM780; Carl Zeiss) and processed using Adobe Photoshop.

For X-gal staining, kidneys/ureters were dissected from non-perfused animals, kidneys were cut in two according to the sagittal plane, and tissues were fixed for 1 h in 1% PFA. Detection of β-galactosidase activity was done on tissues or on 14-μ.m cryostat sections incubated in the dark in staining solution at 37 °C. X-gal staining was performed as described ^38^.

Samples were paraffinized after washing in increasing alcohol concentrations (70%, 80%, 90%, 97% and 100%) VWR chemicals for one day and finally with Xylene 33817 (Sigma-Aldrich) for 2 h, and immersed in paraffin Paraplast X-TRA from Sigma at 65 °C. Staining was performed with Hematoxylin (HHS32-1L Lot. 064K4354 Sigma-Aldrich), Eosin (Ht110230-1L Sigma-Aldrich), Trichrome Masson (Lot.17301-V04 Ral Diagnostics).

### Transmission electronic microscopy and quantification

Kidneys were perfused as described above, rapidly dissected and postfixed in 2% PFA, 2.5% glutaraldehyde in cacodylate buffer (pH 7.2) overnight at 4 °C. The kidneys were put in 1% OsO_4_ solution in cacodylate buffer for 1 h on ice, then dehydrated on ice and embedded in resin (EPON 912). Sample were polymerized 48 h at 60 °C. Ultrathin sections (80-nm) performed on Leica UCT were poststained with 2% uranyl acetate, followed by Reynolds’ lead citrate. Section were examined with a high-resolution transmission electron microscope (Tecnai G2 (FEI), Netherland) at 200 kV and images were acquired with a Veleta camera (Olympus). EM images were opened and analyzed with ImageJ software ^39^; the straight-line tool was used to measure GBM thickness and endothelial cell fenestration on randomly selected electron micrographs. The same approach was used for morphometric analysis of foot process effacement, as described previously ^40^,^41^.

### Blood samples

Blood samples were collected from *Tshz3^+/LacZ^* (n= 11; 5 females and 6 males) and WT mice (n=11; 6 females and 5 males) at 60 days-of-age. The mean values for body weight of *Tshz3^+/LacZ^* and wild-type were 38.27± 7.8 (n=11) and 33.01 ± 4.9 (n=11). Mice were anaesthetized by an intraperitoneal injection of ketamine/xylazine (0.1 ml/10 g body weight) prior to manipulation. Anesthesia was maintained by using 1.7% to 2.5% isoflurane delivered in 600 ml/min oxygen and a closely fitting facemask. Blood was collected by cardiocentesis puncture in heparinize tubes with EDTA and also in tubes without anticoagulant, centrifuging at 4 °C immediately after extraction. The total blood volume in 30-40 g mice is approximately 2 to 3 ml. The maximum volume that could be collected safely at a single survival time point was approximately 800-1000 μl. Blood tests were outsourced to Laboklin G.m.b.H. and performed using a Siemens’ high-volume hematology analyzer (ADVIA 2120i) and a Roche’ chemistry analyzer (Cobas 8000 c701).

### Urine samples

Urine samples were collected from *Tshz3^+/LacZ^* (n= 7; 4 females and 3 males) and WT mice (n=7; 6 females and 1 males) at 60 days-of-age.

Animals were placed in a clean, dry, empty, and transparent individual cage. A non-absorbable plastic, fully sanitizable material was laid on the floor of the cage. During urine collection, water bottles were provided but no food was given to limit contamination with faeces or animal feed. The mouse was monitored all time and removed as soon as it urinated. The voided urine was aspirated with a Gilson Pipetman and transfert to a 1.5 mL sterile microcentrifuge tube (kept on ice). Collected urine was stored at -80C prior to analysis. Due to the small amount of urine collected, the procedure was repeated during three consecutive days to obtain 300ul of urine for each mouse.

### Urinary Proteomics

#### Sample Preparation

Urine samples were collected and frozen at -80°C as described above. Immediately before preparation, mice urine samples were thawed on ice. 150 μl of urine was mixed with a similar volume of a solution containing 2 M urea, 10 mM NH_4_OH, and 0.02% sodium dodecyl sulfate (SDS). To remove high molecular weight molecules, samples were ultrafiltrated (3,400 × g for 45 min at 4°C) using a Centrisart 20kDa cut-off centrifugal filter device (Satorius, Göttingen, Germany) until 200 μl of filtrate was obtained. Afterwards, the filtrate was desalted by a NAP5 gel filtration column (GE Healthcare BioSciences, Uppsala, Sweden) to eliminate electrolytes and urea, hence decreasing matrix effects. The samples were lyophilized in a Savant SpeedVac SVC100H connected to a Virtis 3L Sentry freeze dryer (Fischer Scientific, Illkirch, France), consequently stored at 4°C. Shortly before CE-MS analysis, the samples were resuspended in 10 μl high-performance liquid chromatography grade water (HPLC-H_2_O).

#### CE-MS analysis and data processing

CE-MS experiments were conducted as previously reported ^42^. Briefly, a Beckman Coulter Proteome Lab PA800 capillary electrophoresis system (Fullerton, CA) online coupled to a micrOTOF II MS (Bruker Daltonic, Bremen, Germany) was used. The electro-ionization (ESI) sprayer (Agilent Technologies, Palo Alto, CA) was grounded, and the ion spray interface potential was established to –4.5 kV. Subsequently, data acquisition and MS acquisition methods were automatically measured through the CE by contact-close-relays. Spectra were accrued every 3 seconds, over a range of m/z 350 to 3000.

Mass spectral ion peaks signifying identical molecules at different charges were deconvoluted into singles masses using MosaiquesVisu ^43^. The subsequent peak lists categorized each peptide according to its molecular mass (kDa), CE-migration time (min) and signal intensity (amplitude). Due to the analytical variances of urine samples, migration time and ion signal intensity (amplitude) were normalized using endogenous “housekeeping” peptides, normally displaying a small difference between at least 90% of all urine samples, as reported elsewhere ^42^. Lastly, all detected peptides were deposited, matched and annotated in a Microsoft SQL database ^42^. Thus, further comparisons and statistical analysis among both groups were performed.

#### Sequencing of peptides

Tandem mass spectrometry (MS/MS) analysis were conducted to retrieve the sequence information of the peptides, as previously described ^42^. Briefly, MS/MS experiments were performed on a Dionex Ultimate 3000 RSLC nanoflow system (Dionex, Camberly, UK) coupled to an Orbitrap Velos MS instrument (Thermo Fisher Scientific). Thereafter, all resulting data files were evaluated by the use of SEQUEST (using Thermo Proteome Discoverer 1.2) without any enzyme specificity and searched beside the Swiss-Prot *Mus Musculus* database, as previously described^44^.

#### Statistical analysis

For the identification of potential significant different urinary peptides, urine samples from both groups (wild type and *Tshz3^+/lacZ^*) were compared. *P*-values were calculated according to the Wilcoxon Rank-Sum test. For multiple testing correction, the reported *p*-values were further adjusted via false discovery rate method described by Benjamini and Hochberg ^45^. Only peptides with p-values less than 0.05 and detected in a frequency threshold of ≥70% in at least one of both groups were further considered as statistically significant. Statistical analysis was performed using Prism 7.05 (GraphPad Software, USA) and results considered significant at *P* < 0.05.

## Data availability

The data that support the findings of this study are available from the corresponding author upon reasonable request. Raw data (FastQ files) from the sequencing experiment (triplicates from wild-type and *Tshz3*^+/lacZ^ adult kidney) and raw abundance measurements for genes (read counts) for each sample are available from Gene Expression Omnibus (GEO) under accession GSE182010, which should be quoted in any manuscript discussing the data.

## Ethics Statement

The animal study was reviewed and approved by *“Comité National de Réflexion Ethique sur l’Expérimentation Animale* 14” (ID numbers 57-07112012) and were in agreement with the European Communities Council Directive (2010/63/EU).

## Acknowledgements

We appreciate feedback on the manuscript provided by Pierre L. Roubertoux and Adrian S. Woolf. Microscopy was performed at the imaging platform of the IBDM, supported by the ANR through the “Investments for the Future” program (France-BioImaging, ANR-10-INSB-04-01).

