## Supplementary material for "Haploinsufficiency of the mouse *Tshz3* gene leads to kidney dysfunction": Suplemental figures

### Supplementary Figures

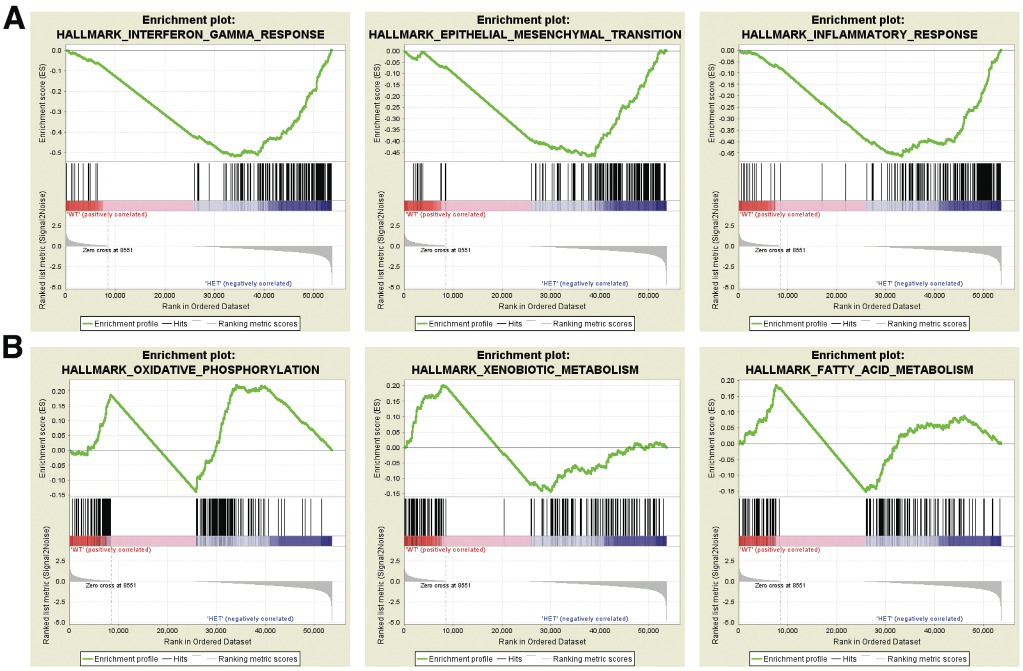


**Supplementary figure 1: Enrichment plots from gene enrichment analysis (GSEA).** GSEA results showing positive **(A)** and negative (**B**) enrichment in the kidneys of *Tshz3*$^{+/lacZ}$ adult mice for gene sets related to "interferon gamma response", "epithelial to mesenchymal transition", "inflammatory response" (**A**) and to "Oxydative Phosphorylation", "Xenobiotic Metabolism" and Fatty Acid Metabolism (**B**).

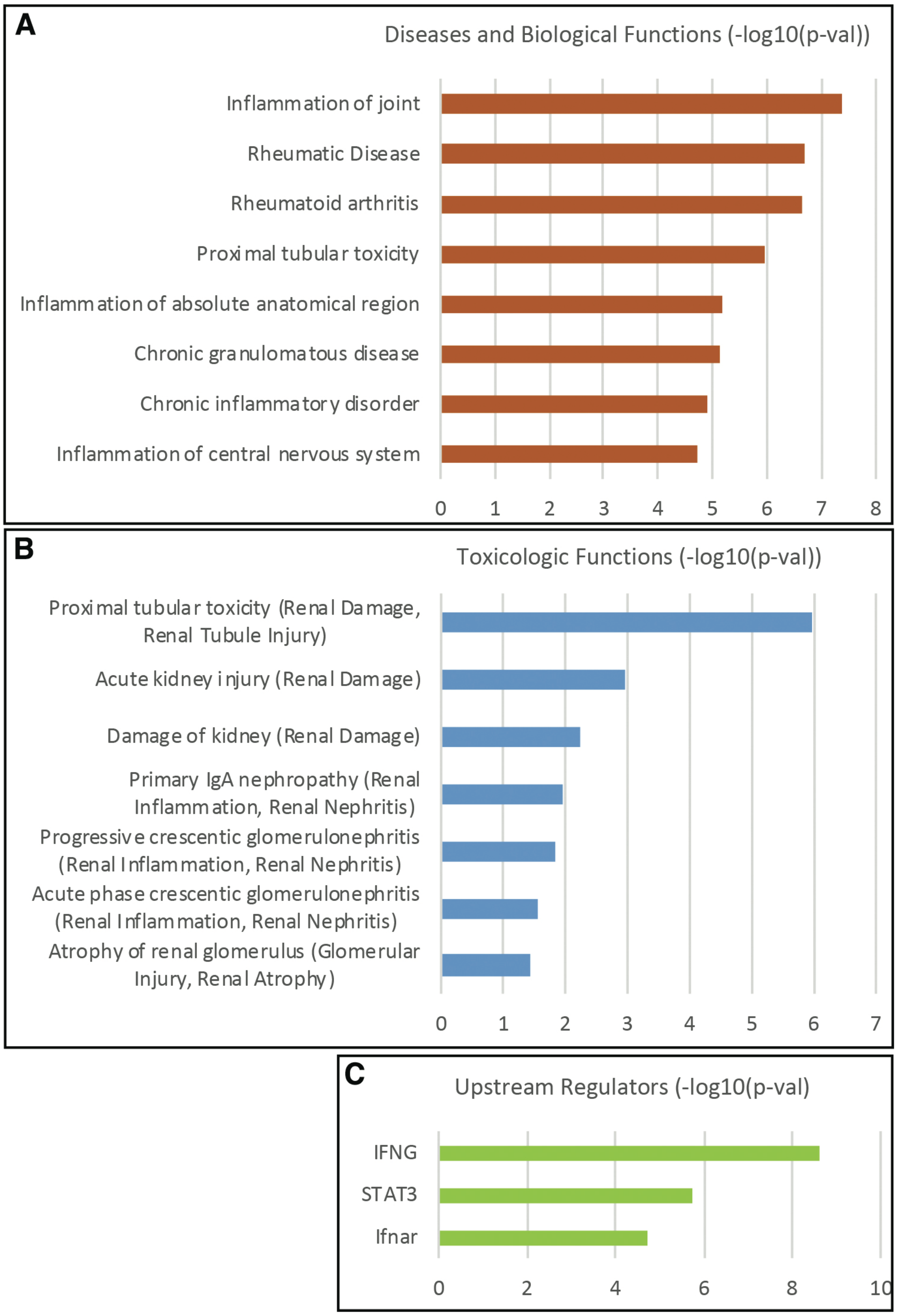


**Supplementary Figure 2: Functional characterization of differentially expressed genes (DEGs) in *Tshz3*$^{+/lacZ}$ adult kidneys, using Ingenuity Pathway Analysis (IPA).** (A) Significant enrichments observed in the Diseases and Biological Functions annotations. (B) Relevant toxicity phenotypes and clinical pathology endpoints associated with the DEGs. (C) Major upstream regulators of the DEGs. Minus Log of the p-values calculated from Fisher's exact test are shown on the x-axis.

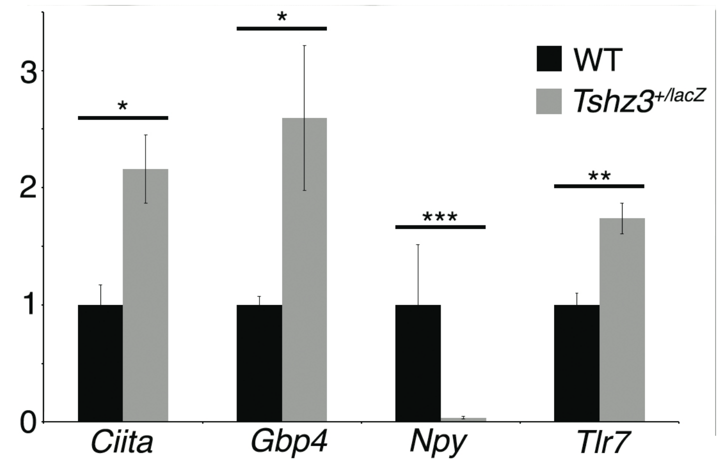


**Supplementary figure 3: Quantitative real-time PCR validation of four differentially expressed genes in *Tshz3*$^{+/lacZ}$ kidney as identified by RNAseq.** Bar graph showing significant variation in mRNA levels as determined by qRT–PCR in *Tshz3*$^{+/lacZ}$ versus WT mice for *Ciita*, *Gbp4*, *Npy* and *Tlr7*. Data are shown as means ± s.e.m. Unpaired two-tailed t test. *, $P$ <0.03; **, $P$ <0.006; ***, $P$ <0.002.

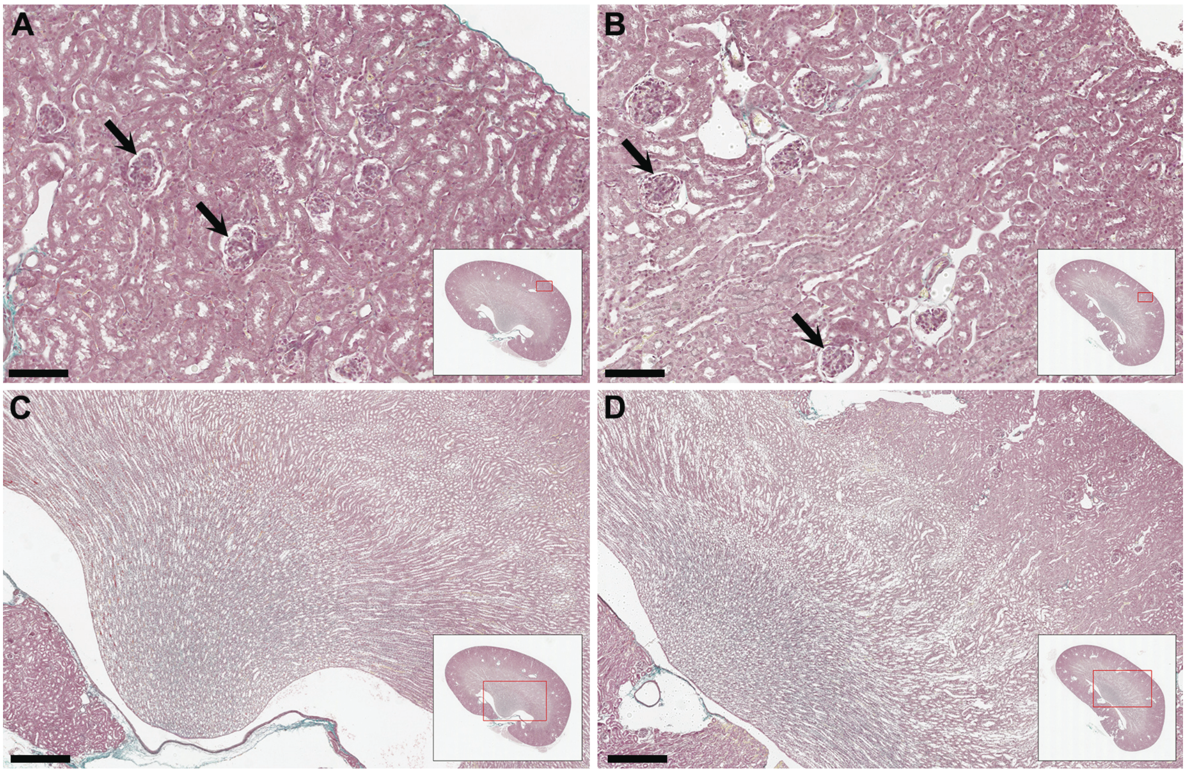


**Supplementary Figure 4. Histological analysis of WT and *Tshz3*$^{+/lacZ}$ kidney.** Representative images of haematoxylin and eosin-stained sections of WT (A, C) and *Tshz3*$^{+/lacZ}$ (B, D) adult kidneys at post-natal day 60. Sections of the cortical region (A, B) and the papilla (C, D). Arrows indicate glomeruli. In the box showing the whole section, the red rectangle indicates which part of the section is shown as A, B, C or D. Scale bars, in A and B, 125µm; in C and D, 500µm.
